# Impact of the X chromosome and sex on regulatory variation

**DOI:** 10.1101/024117

**Authors:** Kimberly R. Kukurba, Princy Parsana, Kevin S. Smith, Zachary Zappala, David A. Knowles, Marie-Julie Favé, Xin Li, Xiaowei Zhu, James B. Potash, Myrna M. Weissman, Jianxin Shi, Anshul Kundaje, Douglas F. Levinson, Philip Awadalla, Sara Mostafavi, Alexis Battle, Stephen B. Montgomery

**Author notes:** Contact information of corresponding authors: Alexis Battle Department of Computer Science, Johns Hopkins University, 3400 N. Charles St., Baltimore, MD 21218, Stephen B. Montgomery Departments of Pathology and Genetics, Stanford University, 300 Pasteur Drive, Stanford, CA 94305.

## Abstract

The X chromosome, with its unique mode of inheritance, contributes to differences between the sexes at a molecular level, including sex-specific gene expression and sex-specific impact of genetic variation. We have conducted an analysis of the impact of both sex and the X chromosome on patterns of gene expression identified through transcriptome sequencing of whole blood from 922 individuals. We identified that genes on the X chromosome are more likely to have sex-specific expression compared to the autosomal genes. Furthermore, we identified a depletion of regulatory variants on the X chromosome, especially among genes under high selective constraint. In contrast, we discovered an enrichment of sex-specific regulatory variants on the X chromosome. To resolve the molecular mechanisms underlying such effects, we generated and connected sex-specific chromatin accessibility to sex-specific expression and regulatory variation. As sex-specific regulatory variants can inform sex differences in genetic disease prevalence, we have integrated our data with genome-wide association study data for multiple immune traits and to identify traits with significant sex biases. Together, our study provides genome-wide insight into how the X chromosome and sex shape human gene regulation and disease.

## Introduction

Many human phenotypes are sexually dimorphic. In addition to males and females having recognizable anatomic and morphological differences, accumulating evidence suggests that they exhibit differences in the prevalence, severity, and age of complex diseases. Classic examples of sex-biased disease include autoimmune disorders (Whitacre et al. 1999; Whitacre 2001), cardiovascular disease (Lerner and Kannel 1986; Mendelsohn and Karas 2005), cancer susceptibility (Cohn et al. 1996; Naugler et al. 2007), and psychiatric disorders (Breslau et al. 1997; Pigott 1999; Hankin and Abramson 2001). While genetic factors may underlie observed differences, determining the genetic contribution to sexual dimorphism has generally lagged behind the hormonal contribution due to challenges in both study design and statistical power (Luan et al. 2001; Patsopoulos et al. 2007; Ober et al. 2008). Despite these limitations, several studies have discovered genotype-by-sex interaction effects in human phenotypes, such as anthropometric traits (Heid et al. 2010; Randall et al. 2013), bone mineral density (Liu et al. 2012a), complex diseases (Liu et al. 2012b; Myers et al. 2014), and the intermediate phenotype of gene expression (Dimas et al. 2012b; Yao et al. 2014). To explain the etiology of these sexually dimorphic traits, several mechanisms have been proposed, including those arising due to the X chromosome (Dobyns et al. 2004; Ober et al. 2008).

Although genome-wide association studies (GWAS) have uncovered numerous loci associated with complex phenotypes on the autosomes, the X chromosome is significantly underrepresented in such work. Indeed, only one-third of GWAS include the X chromosome, largely due to specialized analytical methods required for processing and interpreting genetic data on this chromosome (Wise et al. 2013). Furthermore, many large-scale functional genomic studies investigating the effect of genetic variants also exclude the X chromosome (Dimas et al. 2009; Montgomery et al. 2010; Pickrell et al. 2010; Lappalainen et al. 2013; Battle et al. 2014; Consortium 2015). Motivated by the underutilization of the X chromosome, recent studies have characterized the role of the X chromosome in the heritability of human phenotypes (Chang et al. 2014; Tukiainen et al. 2014). However, few studies have systematically investigated the contribution of the X chromosome in the context of both regulatory variation and its interaction with sex.

In this study, we report on the impact of sex and the X chromosome on human gene expression as an intermediate phenotype to help understand the genetic and molecular basis of sex-biased disease risk. First, we examine differential gene expression between the sexes and evaluate the proportion of variation explained by genotype. Second, as one of the first and largest studies to our knowledge to test for expression quantitative trait loci (eQTL) on the X chromosome, we observe a depletion of eQTL relative to autosomes, suggestive of more efficient purifying selection on the X chromosome. In contrast, we detect an enrichment of sex-interacting eQTLs on the X chromosome. To identify the underlying mechanisms of sex-specific expression and genotype-by-sex interactions, we generated chromatin accessibility data and tested the relationship between sex-interacting eQTL and sex-specific chromatin accessibility patterns. Lastly, we evaluate the sex-specific effects of genotype on expression for trait-associated loci identified in genome-wide association studies (GWAS) and observe sex-biased effects for multiple immune-associated diseases. In summary, our investigation into sex-specific gene expression and genetic architecture provides new insight into the regulatory mechanisms of sexual dimorphism in human phenotypes and the importance of including the X chromosome and sex in the design, analysis, and interpretation of genetic studies.

## Results

To study sex-specific genetic variation in humans, we obtained gene expression data for the Depression Genes and Networks (DGN) cohort comprised of 922 individuals of European ancestry across the United States (Battle et al. 2014; Mostafavi et al. 2014). Gene and isoform expression was quantified from whole blood 51-bp single-end RNA-sequencing data (Methods). Each individual was also genotyped for 737,187 single nucleotide polymorphisms (SNPs) located on the autosomes and X chromosome on the Illumina HumanOmni1-Quad BeadChip. We then imputed variants using the 1000 Genomes Phase 1 reference panel (Methods).

### Sex-specific expression variation across the genome

Beyond differences in mean expression levels, differences in expression variation may be attributed to interaction effects of genomic variants and both environmental and biological variables, such as sex (Idaghdour and Awadalla 2012). We investigated sex-specific patterns of gene expression and variance across the genome. After correcting for known technical covariates in the expression data (see Methods), we tested if gene expression variance in females is equivalent to that in males (using an F-test). We limited our analysis to genes expressed in both sexes and matched the number of males and females tested to ensure that sample size did not influence measured variance (Methods). We identified 487 genes with sex-specific expression variance (5% FDR). Across the genome, we observed that a higher proportion of genes with sex-specific variance are on the X chromosome compared to autosomes (9.8% versus 6.4%, *P*-value = 1.9 × 10^−3^, Fisher’s exact test; Fig. 1A). This pattern replicates in the ImmVar cohort providing cell-type specific data (Ye et al. 2014) (*P*-value < 5.4 × 10^−3^; Fig. S1), indicating that this is unlikely to be an artifact of differences in cell type proportions between sexes. As with variance, we observe more genes on the X chromosome exhibiting sex-specific gene expression compared to autosomes (54.8% versus 48.4%, *P*-value = 4.4 × 10^−3^, Fisher’s exact test; Fig. 1A), an observation predicted by theory (Rice 1984) and seen in other species (Ranz et al. 2003; Yang et al. 2006). To account for any potential bias in the relationship of variance and mean, we tested whether genes with sex-specific expression variance are genes with a significant difference in their mean expression between the sexes (FDR < 0.05, Welch’s two-sample *t*-test) and detected no significant enrichment (*P*-value = 0.39, chi-square test). Furthermore, since variance may increase as the mean increases for RNA-sequencing data, we tested if genes with higher variance in one sex also show higher expression and observed that genes with higher variance are more likely to have lower expression (47.8% versus 52.2%, *P*-value = 1.1 × 10^−7^, binomial exact).

**Figure 1.**
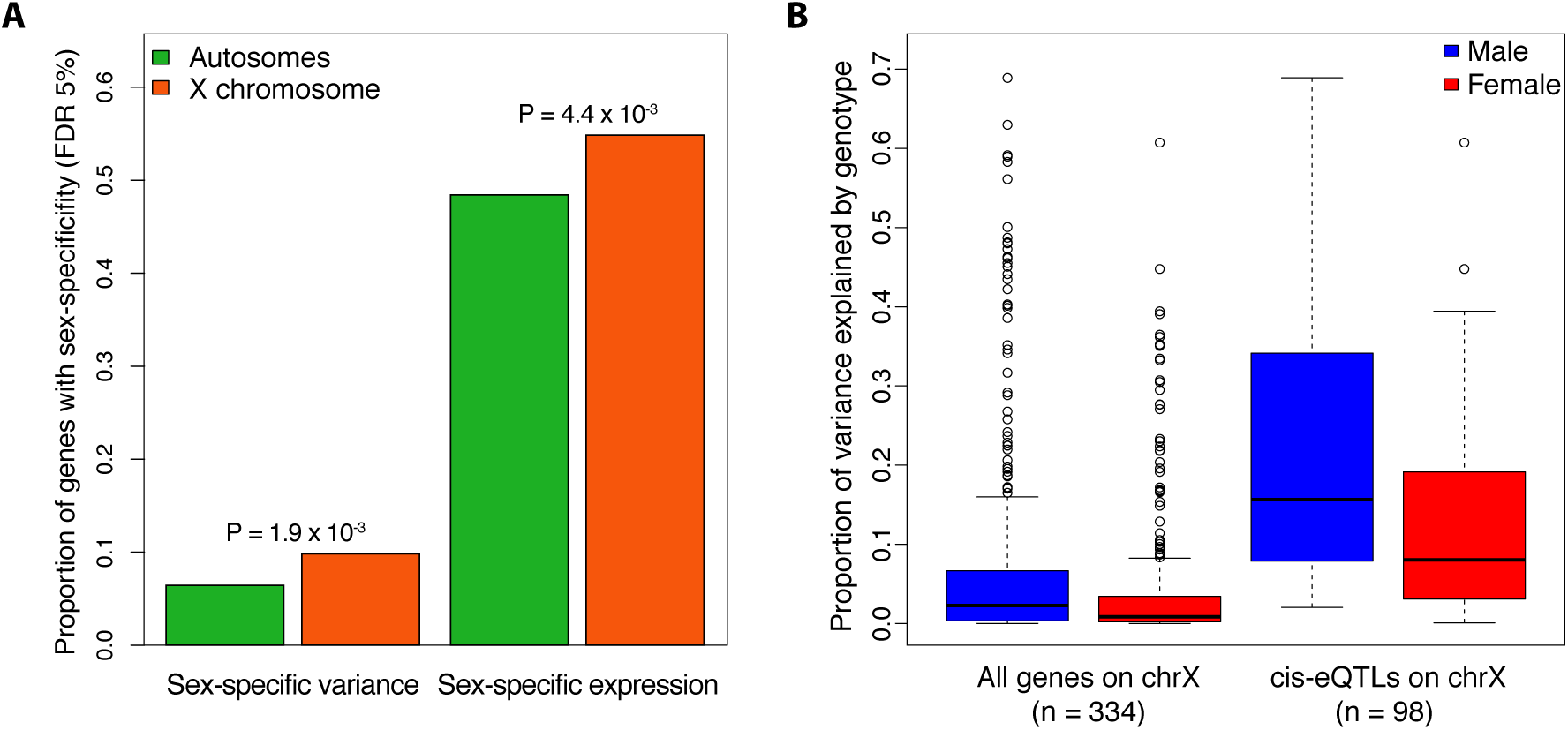
Differential expression variance within the sexes. **A.** Comparison of genes with significant sex-specific expression variance (FDR 5%) and sex-specific expression (FDR 5%) on the autosomes and the X chromosome in the DGN. To test for differences in mean expression and variance, the number of males and females were matched (n = 274). One-sided Fisher’s exact test *P*-values above bars indicate significance of higher expression or variance on X chromosome relative to autosomes. **B**. Proportion of variance explained (PVE) by genotype on the X chromosome in males and females. To test for the PVE, the number of males and females were matched (n = 274). We tested all genes on the X chromosome and genes with a *cis-*eQTL (Bonferroni adjusted *P*-value < 0.05)

To understand why the X chromosome exhibits unique patterns of expression variance relative to the autosomes, we tested if the hemizygosity of the X chromosome contributes to greater expression variance in males. To test this hypothesis, we estimated the proportion of expression variance explained by common genetic variants (MAF > 0.05) in males and females for each gene on the X-chromosome. We found that the mean estimated percentage of variance explained by genotype is 1.7-fold higher for males than females (2.3% versus 0.9%; *P*-value = 6.14 × 10^−5^; Wilcoxon-rank-sum test; Fig. 1B). This difference remained when restricting only to genes with a *cis*-eQTL (Bonferroni-adjusted *P*-value < 0.01; 21.0% versus 12.5%; *P*-value = 6.4 × 10^−5^; Wilcoxon-rank-sum test; Fig. 1B). In contrast, if we evaluate the proportion of variance explained by genotype on an autosomal chromosome, we observe no significant difference between males and females (*P*-value = 0.557 and 0.686, Wilcoxon-rank-sum test; Fig. S2). We then asked if the autosomal genes with sex-specific expression variance are enriched in specific biological processes and found that these genes are enriched in cell death and regulation of apoptosis (Table 1). These observations suggest that divergent regulation of cell death occurs between the sexes, supporting previous studies that have found sex-specific alterations in apoptosis within immune and neural cell populations (Molloy et al. 2003; Lang and McCullough 2008).

**Table 1.** Functional enrichment of genes with sex-specific expression variance (FDR 5%)

| Category | Term | Count | % | P-Value | Fold Enrichm | Fisher Exact |
| --- | --- | --- | --- | --- | --- | --- |
| GOTERM_BP_FAT | positive regulation of apoptosis | 7 | 18.9 | 4.40E-05 | 9.4 | 4.60E-06 |
| GOTERM_BP_FAT | positive regulation of programmed cell death | 7 | 18.9 | 4.60E-05 | 9.3 | 4.80E-06 |
| GOTERM_BP_FAT | positive regulation of cell death | 7 | 18.9 | 4.70E-05 | 9.3 | 5.00E-06 |
| GOTERM_BP_FAT | induction of apoptosis | 6 | 16.2 | 1.40E-04 | 10.6 | 1.30E-05 |
| GOTERM_BP_FAT | induction of programmed cell death | 6 | 16.2 | 1.40E-04 | 10.6 | 1.30E-05 |
| GOTERM_BP_FAT | regulation of small GTPase mediated signal transduction | 5 | 13.5 | 7.10E-04 | 11.2 | 6.20E-05 |
| GOTERM_BP_FAT | induction of apoptosis by extracellular signals | 4 | 10.8 | 8.50E-04 | 19.8 | 4.20E-05 |
| GOTERM_BP_FAT | regulation of apoptosis | 7 | 18.9 | 1.10E-03 | 5.2 | 2.00E-04 |
| GOTERM_BP_FAT | regulation of programmed cell death | 7 | 18.9 | 1.10E-03 | 5.2 | 2.10E-04 |
| GOTERM_BP_FAT | regulation of cell death | 7 | 18.9 | 1.20E-03 | 5.1 | 2.20E-04 |
| GOTERM_BP_FAT | apoptosis | 6 | 16.2 | 2.10E-03 | 5.9 | 3.50E-04 |
| GOTERM_BP_FAT | programmed cell death | 6 | 16.2 | 2.20E-03 | 5.8 | 3.70E-04 |
| GOTERM_BP_FAT | membrane organization | 5 | 13.5 | 2.60E-03 | 7.8 | 3.30E-04 |
| GOTERM_BP_FAT | cell death | 6 | 16.2 | 4.30E-03 | 5 | 8.30E-04 |
| GOTERM_BP_FAT | death | 6 | 16.2 | 4.40E-03 | 5 | 8.60E-04 |
| GOTERM_BP_FAT | regulation of Ras protein signal transduction | 4 | 10.8 | 4.70E-03 | 10.9 | 4.20E-04 |
| GOTERM_BP_FAT | regulation of Rho protein signal transduction | 3 | 8.1 | 1.10E-02 | 17.7 | 6.00E-04 |
| GOTERM_BP_FAT | mitotic cell cycle | 4 | 10.8 | 2.50E-02 | 5.9 | 4.10E-03 |
| GOTERM_BP_FAT | cellular protein localization | 4 | 10.8 | 2.90E-02 | 5.6 | 4.90E-03 |
| GOTERM_BP_FAT | cellular macromolecule localization | 4 | 10.8 | 2.90E-02 | 5.5 | 5.10E-03 |
| GOTERM_BP_FAT | membrane invagination | 3 | 8.1 | 4.70E-02 | 8.2 | 5.50E-03 |
| GOTERM_BP_FAT | endocytosis | 3 | 8.1 | 4.70E-02 | 8.2 | 5.50E-03 |
| GOTERM_BP_FAT | actin cytoskeleton organization | 3 | 8.1 | 4.80E-02 | 8 | 5.90E-03 |
| GOTERM_BP_FAT | actin filament-based process | 3 | 8.1 | 5.30E-02 | 7.6 | 6.80E-03 |
| GOTERM_BP_FAT | vesicle-mediated transport | 4 | 10.8 | 6.10E-02 | 4.1 | 1.40E-02 |
| GOTERM_BP_FAT | cell cycle process | 4 | 10.8 | 6.60E-02 | 4 | 1.60E-02 |
| GOTERM_BP_FAT | cell morphogenesis | 3 | 8.1 | 8.10E-02 | 5.9 | 1.30E-02 |
| GOTERM_BP_FAT | regulation of hydrolase activity | 3 | 8.1 | 8.40E-02 | 5.8 | 1.40E-02 |
| GOTERM_BP_FAT | cell division | 3 | 8.1 | 8.50E-02 | 5.8 | 1.40E-02 |
| GOTERM_BP_FAT | cell projection organization | 3 | 8.1 | 9.20E-02 | 5.5 | 1.60E-02 |
| GOTERM_BP_FAT | cellular component morphogenesis | 3 | 8.1 | 9.80E-02 | 5.3 | 1.80E-02 |
| GOTERM_BP_FAT | cell cycle | 4 | 10.8 | 1.30E-01 | 3 | 4.10E-02 |
| GOTERM_BP_FAT | cytoskeleton organization | 3 | 8.1 | 1.50E-01 | 4.2 | 3.30E-02 |
| GOTERM_BP_FAT | cell cycle phase | 3 | 8.1 | 1.50E-01 | 4.2 | 3.40E-02 |
| GOTERM_BP_FAT | protein localization | 4 | 10.8 | 1.70E-01 | 2.6 | 6.00E-02 |
| GOTERM_BP_FAT | proteolysis | 4 | 10.8 | 1.80E-01 | 2.5 | 6.70E-02 |
| GOTERM_BP_FAT | immune response | 3 | 8.1 | 2.30E-01 | 3.1 | 6.80E-02 |
| GOTERM_BP_FAT | modification-dependent protein catabolic process | 3 | 8.1 | 2.40E-01 | 3.1 | 7.20E-02 |
| GOTERM_BP_FAT | modification-dependent macromolecule catabolic process | 3 | 8.1 | 2.40E-01 | 3.1 | 7.20E-02 |
| GOTERM_BP_FAT | proteolysis involved in cellular protein catabolic process | 3 | 8.1 | 2.60E-01 | 2.9 | 8.10E-02 |
| GOTERM_BP_FAT | cellular protein catabolic process | 3 | 8.1 | 2.60E-01 | 2.9 | 8.10E-02 |
| GOTERM_BP_FAT | protein catabolic process | 3 | 8.1 | 2.70E-01 | 2.8 | 8.70E-02 |
| GOTERM_BP_FAT | intracellular signaling cascade | 4 | 10.8 | 2.90E-01 | 2 | 1.30E-01 |
| GOTERM_BP_FAT | intracellular transport | 3 | 8.1 | 2.90E-01 | 2.6 | 1.00E-01 |
| GOTERM_BP_FAT | cellular macromolecule catabolic process | 3 | 8.1 | 3.30E-01 | 2.4 | 1.20E-01 |
| GOTERM_BP_FAT | macromolecule catabolic process | 3 | 8.1 | 3.70E-01 | 2.2 | 1.50E-01 |
| GOTERM_BP_FAT | regulation of transcription | 4 | 10.8 | 7.30E-01 | 1 | 5.50E-01 |

### **Identification and characterization of X-chromosome *cis-*eQTLs**

We tested for genetic variants that affect local gene expression on the autosomes and X chromosome. For each gene, we evaluated the association between overall gene expression and genetic variants within 1 Mb of the transcription start site (TSS) using linear regression. At a genome-wide level FDR of 5%, we detect eQTLs on 74.8% of autosomal genes and 43.7% of X-chromosome genes. For a range of FDR thresholds, we find a depletion of eQTLs on the X chromosome relative to autosomes (two-sided chi-square test *P*-value = 9.4 × 10^−37^ at FDR 1%; Fig. 2A). To determine if either male hemizygosity of the X chromosome or female X-inactivation influenced this observed depletion, we also detected eQTLs in just the male and female populations. In both independent male and female populations, we also observed a depletion of eQTLs on the X chromosome across multiple FDR thresholds (*P*-value < 10^−15^, chi-square test; Fig. 2A), compared to autosomal eQTL rates. The depletion of X chromosome eQTLs further replicated in multiple individual cell types, including monocytes from the ImmVar cohort and lymphoblastoid cells from the Geuvadis cohort (Table S1).

**Figure 2.**
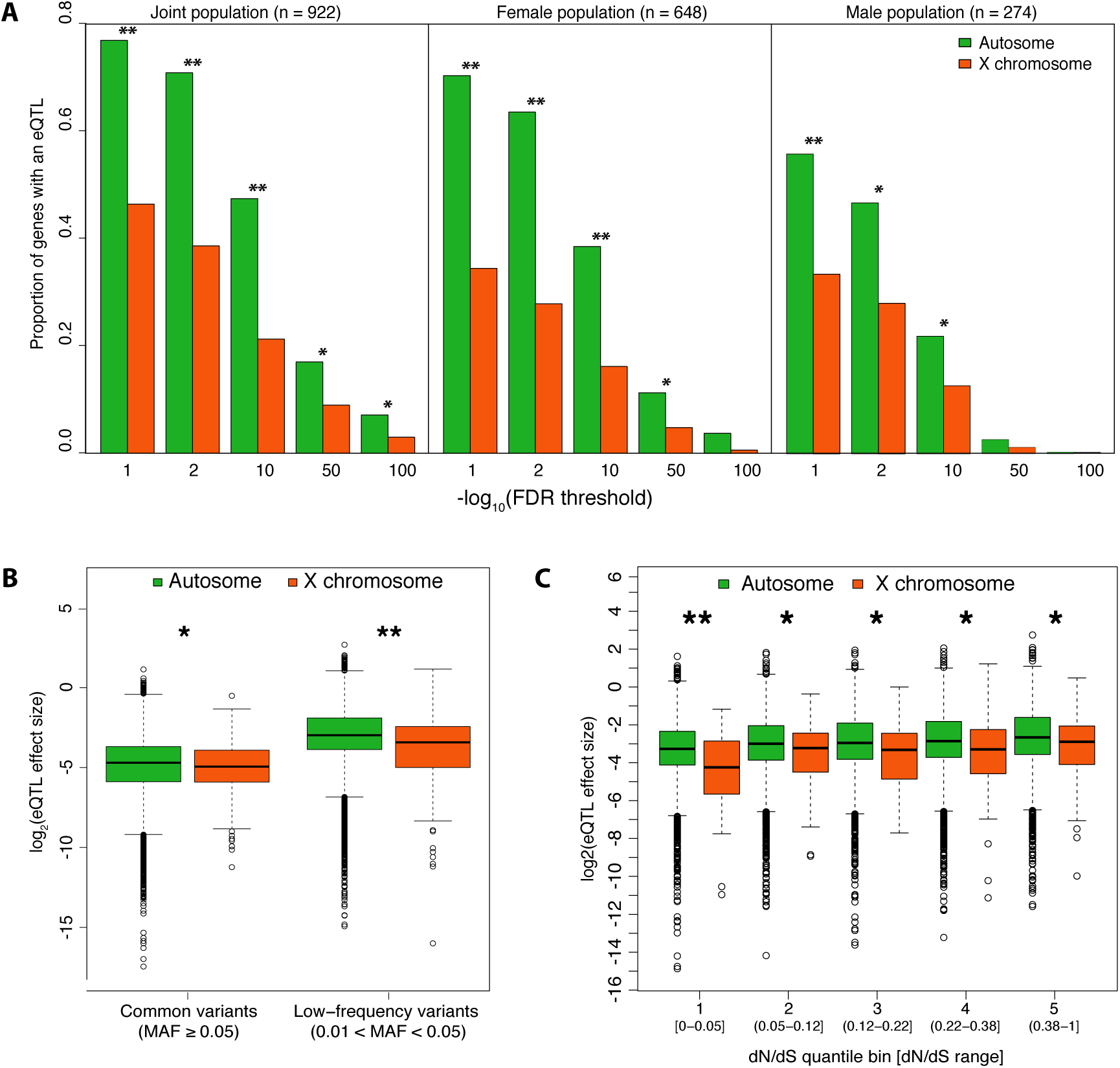
Characterization of eQTLs on the X chromosome. **A.** Proportion of genes with an eQTL at different FDR thresholds discovered in the joint (n = 922), female (n = 648), and male (n = 274) populations. \*\**P*-value < 1 × 10^−15^, \**P*-value < 0.05, Bonferroni adjusted chi-square test. **B**. Comparison of eQTL effect size between autosomes and the X chromosome for common variants (MAF > 0.05) and low-frequency variants (0.01 < MAF< 0.03). The difference in effect size is statistically significant for both common (*P*-value = 0.0382, two-sided Wilcoxon rank sum test) and low-frequency variants (*P*-value = 1.21 × 10^−7^). **C.** Relationship of eQTL effect size and strength of purifying selection. Boxplots for the effect size of eQTLs on the autosomes (n = 11967) and the X chromosome (n = 303) at genes five dN/dS bins. eQTLs on the X chromosome have lower effect sizes compared to eQTLs on the autosomes across different dN/dS bins (one-sided Wilcoxon-rank-sum test; \*\**P*-value < 5 × 10^−4^; * *P-*value < 5 × 10^−2^)

#### eQTL effect sizes on the X chromosome.

To understand the properties of X chromosome eQTLs, we compared the magnitude of the genotypic effect on expression, or the effect size, to that of autosomal eQTLs within females. For eQTLs detected from common variants (MAF ≥ 0.05), we observe that the effect size on the X chromosome is 1.17-fold lower relative to autosomes (*P*-value = 3.8 × 10^−2^; Wilcoxon rank-sum test; Fig. 2B). For eQTLs detected from low-frequency variants (0.01 < MAF < 0.05), we observe that effect size on the X chromosome is 1.34-fold lower relative to autosomes (*P*-value = 1.2 × 10^−7^, Wilcoxon rank-sum test) suggesting increased purifying selection for large eQTL effects on the X chromosome versus the autosomes.

Previous studies have found that the hemizygosity in males may cause unusual patterns of evolution, including lower nucleotide diversity, lower effective population size, and more efficient removal of deleterious alleles on the X chromosome compared to the autosomes (Andolfatto 2001; Schaffner 2004; Lu and Wu 2005; Vicoso and Charlesworth 2006). To test the relationship of eQTL effect size, selective constraint and purifying selection on the X chromosome, we obtained the ratios of non-synonymous to synonymous substitutions (dN/dS) of human-rhesus genes as an indicator of selective constraint at the sequence level, quantile binned the dN/dS ratios and compared the eQTL effect sizes on the X chromosome to autosomes. For genes under the greatest degree of constraint (lower 20% quantile bin, dN/dS 0-0.05), we observe significantly lower effect sizes on the X chromosome (*P*-value = 3.6 × 10^−4^, Wilcoxon rank-sum test; Fig. 2C). For genes under less constraint (higher dN/dS ratios), we observe similar but less significant patterns (*P*-value < 5.0 × 10^−2^). This evidence suggests that even by comparing genes with similar selective constraints, X-chromosome genes experience greater purifying selection on regulatory variation relative to the autosomes.

### Identification and characterization of sex-interacting eQTL

To identify sex-interacting eQTLs, we used a linear model with a genotype-sex interaction term. Genotype-by-sex interactions occur when the effect of genotype on expression differs between males and females. For example the genetic effects on expression may be present in only one sex, may have different magnitudes of effect, or may even have opposing directions of effect in the two sexes (Fig. S3). For each gene, we tested the genotype-sex interaction term for all variants within 1 Mb of the gene. We observe an enrichment of sex-specific eQTLs on the X chromosome compared to the autosomes (Fig. 3A; Wilcox rank-sum test *P*-value = 8.2 × 10^−4^ and 1.9 × 10^−5^ for the top 50 and 500 associations, respectively). After adjusting for the number of SNPs tested per gene using Bonferroni and subsequently identifying sex-interacting eQTLs using gene-level significance at FDR 5%, we discover six eQTLs (four autosomal and two X-chromosome eQTLs) with significant sex interactions (Table 2). If we restrict our tests to variants previously detected as *cis-*eQTLs (Bonferroni adjusted *P*-value < 0.1), we improve power but the total number of discoveries is unaffected (Fig. S4). Notably, two of the sex-interacting eQTLs discovered in our DGN cohort replicate in other studies; rs4785448–*NOD2* and chr11:62735958:D-*BSCL2* replicate in the Framingham Heart Study (Yao et al. 2014) and CARTaGENE Study (Awadalla et al. 2013; Hussin et al. 2015), respectively (Table S2). We additionally used a binomial generalized linear mixed model (Knowles et al., in preparation) to test for a sex-interacting allele-specific expression QTL (aseQTL). Specifically, we tested if alleles at heterozygous loci show allele-specific expression (ASE) and if the magnitude of ASE effects differs in males and females. Two of our six significant sex-interacting eQTL genes had heterozygous loci with sufficient read depth to measure allelic imbalance, and of these two, we observed that *NOD2* has a significant sex-interacting aseQTL (*P*-value = 2.5 × 10^−3^; Fig. S5). All variants tested for sex interactions in the DGN cohort can be browsed at http://montgomerylab.stanford.edu/resources.html.

**Table 2.**
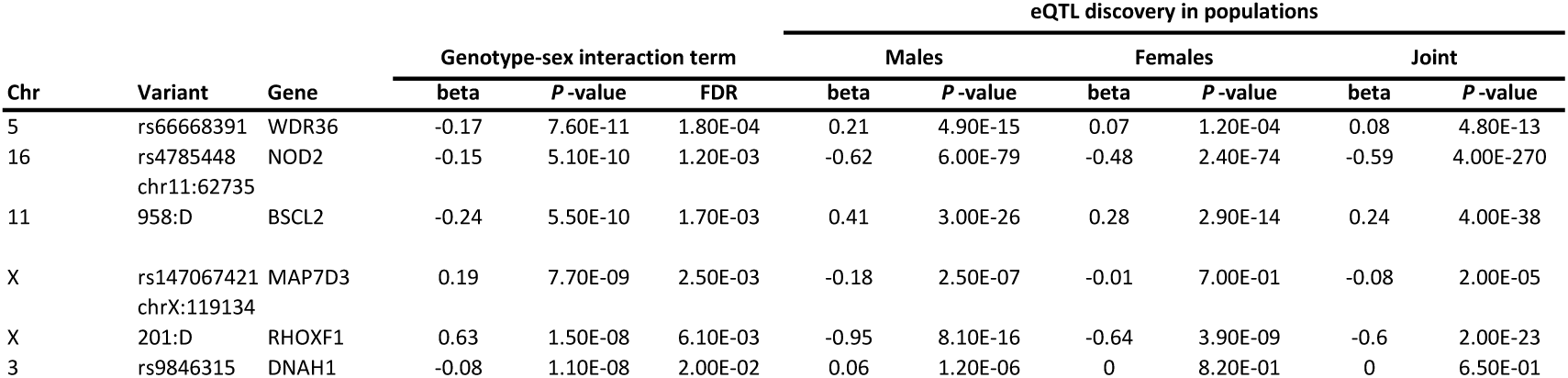
Discovery of sex-interacting eQTLs (FDR 5%)

**Figure 3.**
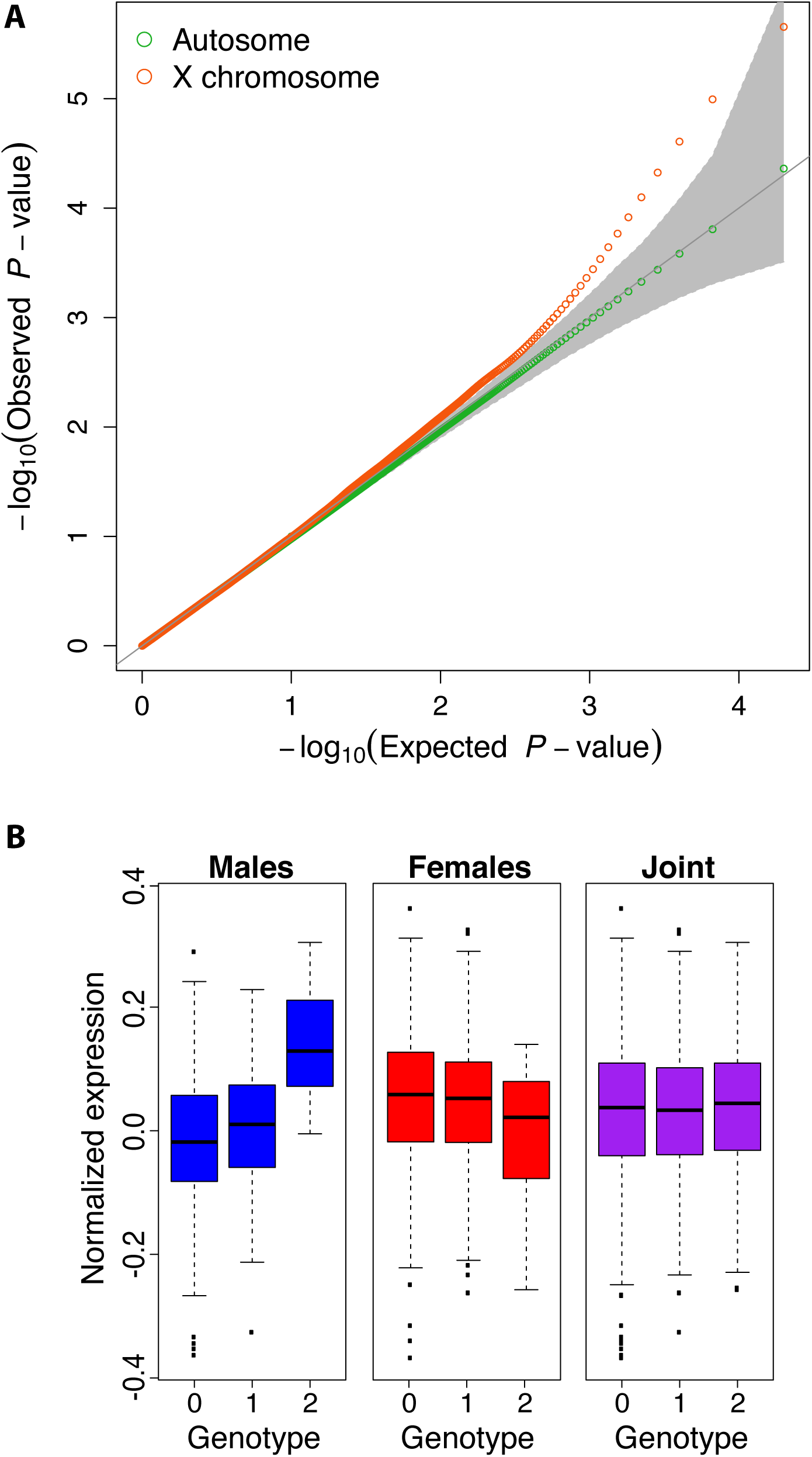
Discovery of sex-interacting eQTLs. **A.** Quantile-quantile (QQ) plot describing the sex-interacting eQTL association P-values for SNPs tested within 1MB of genes on the X chromosome (red) and the autosomes (purple) with 95% confidence interval (gray). **B.** Sex-interacting eQTL (q-value = 0.0198) for *DNAH1* (dynein, axonemal, heavy chain 1), a protein-coding gene involved in microtubule motor activity, ATPase activity, and sperm motility.

Of the six genes with sex-interacting eQTLs, *DNAH1* on chromosome 3 was distinct because this SNP-genotype pair was not a significant eQTL in the female or joint population analysis (Fig. 3B). Although this gene exhibits similar expression between the sexes in whole blood (nominal negative binomial *P*-value = 0.81), it shows the highest expression in the testis compared to the remaining 46 tissues tested in the GTEx Project (Consortium 2013). We investigated this gene further and found that it is a force-producing protein with ATPase activity involved in sperm motility and flagellar assembly (UniProt ID Q9P2D7). The sex-specific function of this gene prompted us to test if the sex-interacting eQTLs were enriched in sex-specific biological processes. To evaluate this hypothesis, we tested the top 100 sex-interacting eQTLs for enrichment in genes involved in sex differentiation (GO:0007548) and discovered a modest enrichment (odds ratio = 6.6 [1.17-18.2], *P*-value = 4.3 × 10^−3^, Fisher’s exact test). When testing all biological processes, no specific process passed multiple testing corrections, however the top GO terms were linked to sex-specific biological processes (e.g. reproductive structure development and response to estrogen stimulus; Table S3).

To determine if sex-interacting eQTL discovery is driven by differences in gene expression between males and females, we tested for enrichment of differentially expressed genes (FDR 5%). We observed no significant enrichment of genes with sex-specific expression for the top 100 sex-interacting eQTLs (odds ratio = 0.4 [0.05-1.58], *P*-value = 3.3 × 10^−1^, Fisher’s exact test) compared to background of genes expressed in whole blood. Concordant with a previous study (Dimas et al. 2012a), this demonstrates that sex-interacting eQTLs likely do not arise as a consequence of expression differences between the sexes and may result from other factors that differ in a sex-specific matter, such as transcription factor activity, hormone receptors, and chromatin accessibility.

### Sex-specific chromatin accessibility

To identify the molecular mechanisms of sex-specific gene regulation, we generated and investigated differences in chromatin accessibility between males and females. We measured chromatin accessibility of peripheral blood mononuclear cells (PBMCs) from twenty individuals matched for age, ethnicity, and sex using the assay for transposase-accessible chromatin followed by sequencing (ATAC-seq) (Buenrostro et al. 2013). Using a negative binomial model, we identified 577 (0.69%) sex-specific chromatin accessibility regions at FDR 5% (see Methods).

Considering the unique heterochromatic state of the X chromosome in females, we asked if the distribution of sex-specific open chromatin regions differed between the autosomes and the X chromosome (Fig. 4A). We observed an enrichment of sex-specific open chromatin regions on the X (Wilcox rank-sum test *P*-value = 3.5 × 10^−10^ and 1.4 × 10^−4^ for the top 50 and 500 associations, respectively). Following this observation, we hypothesized that regions on the X chromosome are more likely to have greater chromatin accessibility in females due to genes escaping X-inactivation. Indeed, we found that X-chromosome regions are more likely to have higher chromatin accessibility in females than males compared to autosomal regions (odds ratio = 2.44 [1.99-3.01], *P*-value < 2.2 × 10^−16^, Fisher’s exact test) (Fig. S6). To further interpret this, we then tested if the top sex-specific peaks on the X chromosome are more likely to be genes escaping X-inactivation (Carrel and Willard 2005; Park et al. 2010). Indeed, we observed that the top 10% of sex-specific peaks on the X chromosome are enriched for genes escaping X-inactivation compared to genes in sex-shared peaks (44.6% versus 6.5%; odds ratio = 9.60 [2.70-34.49], *P*-value = 1.6 × 10^−4^, Fisher’s exact test), indicating that a large number of the X chromosome sex-specific peaks inform regions of escape from X inactivation.

**Figure 4.**
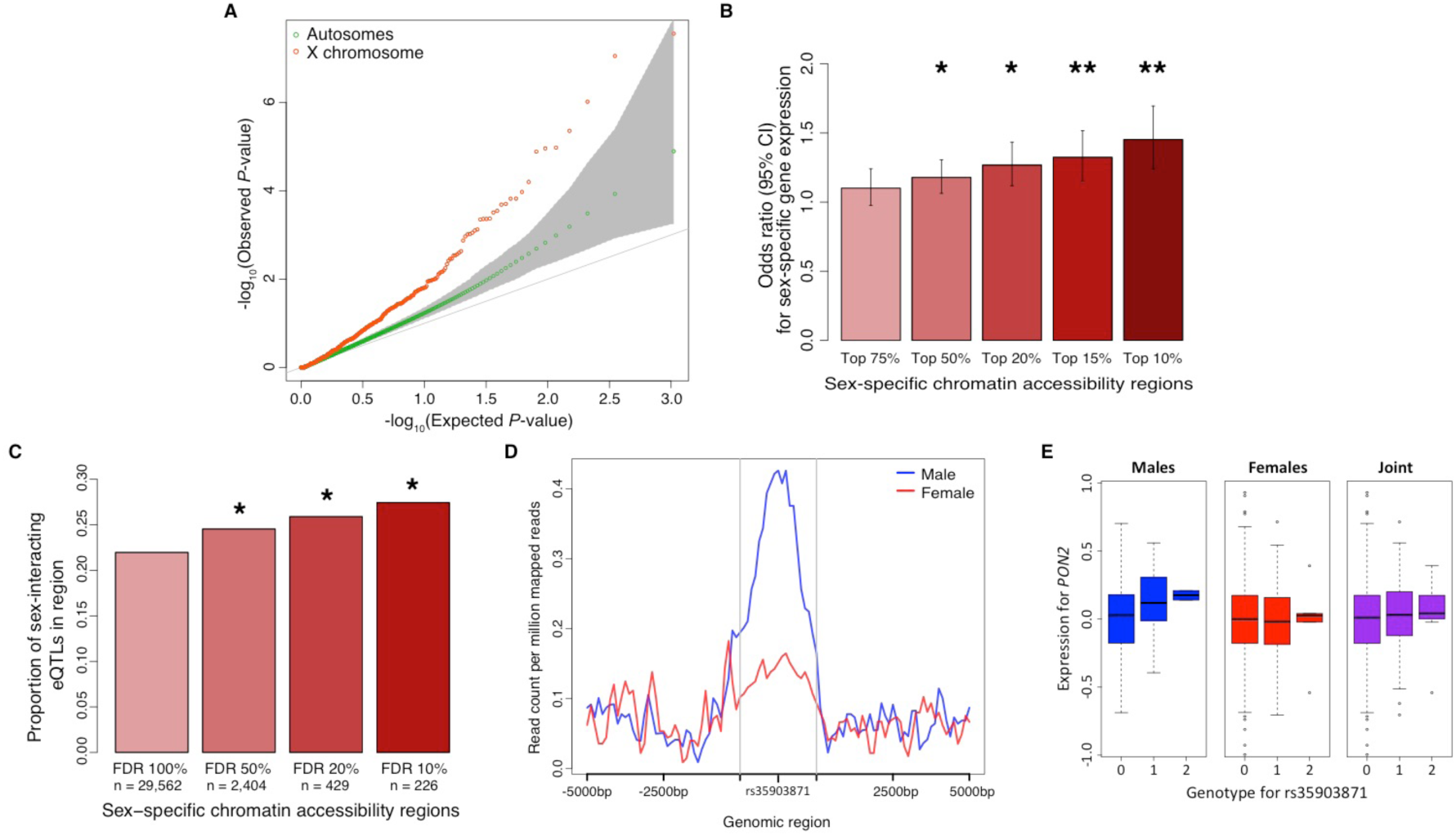
Discovery of sex-specific chromatin accessibility regions. **A.** Q-Q plot for tests of differential chromatin accessibility between the sexes. 95% genome-wide confidence interval in gray **B.** Enrichment of genes with differential expression between the sexes (FDR 5%) with differential chromatin accessibility (varying thresholds) 40kb upstream. \*\**P*-value < 10^−5^, \**P*-value < 10^−3^, Fisher’s exact test. **C.** Proportion of sex-interacting eQTL genes (*P-*value < 0.05) in differential chromatin accessibility regions (varying thresholds). *P-value < 5.0 × 10^−2^, Fisher’s exact test). **D**. Chromatin accessibility peak located at chr7:95,063,722-95,064,222 within 5,000 upstream and downstream. This region has differential chromatin accessibility between males and females (nominal *P-*value = 4.1 × 10^−4^, *Q*-value = 7.5 × 10^−2^). **E.** Sex-interacting eQTL for *PON2* and rs35903871 located at chr7:95,063,972 (nominal *P*-value = 8.0 × 10^−3^)

#### Integration of sex-specific chromatin accessibility with expression.

One intuitive mechanism for sex-specific gene expression and sex-interacting eQTLs is sex-specific chromatin accessibility. Therefore, we sought to identify if differential open chromatin between the sexes was associated with sex-specific expression and genotype-sex interactions. First, we tested if genes with sex-specific expression (FDR 5%) are enriched in regions with sex-specific chromatin accessibility. We observed that genes with sex-specific chromatin accessibility 40 kilobases upstream of the gene TSS were associated with sex-specific expression (odds ratio = 1.45 [1.24-1.69], *P*-value = 4.7 × 10^−6^, Fisher’s exact test) (Fig. 4B).

Next, we investigated if sex-specific chromatin accessibility regions are enriched in sex-interacting eQTL genes by testing for genotype-sex interactions for variants within 1 Mb of the gene. We observe that genes with a sex-interacting eQTL (nominal *P*-value < 0.1) are likely to have differential chromatin accessibility between the sexes (odds ratio = 1.36 [0.99-1.81], *P*-value = 5.1 × 10^−2^, Fisher’s exact test) (Fig. 4C). To ensure that we did not observe this enrichment by chance, we permuted the sexes for eQTL testing and tested sex-interacting eQTLs in sex-specific chromatin regions and observed no enrichment (Fig. S7). In addition, we asked if sex-specific chromatin accessibility regions were enriched in sex-interacting eQTL. We tested the genotype-sex interaction term for the variant closest to the region mid-point and the closest five genes and observed weak but not statistically significant enrichment (Fig. S8). One example of a sex-interacting eQTL in a sex-specific chromatin accessibility region is illustrated in Fig. 4D, in which a SNP (rs35903871) is located in a region with greater chromatin accessibility in males (*P*-value = 4.1 × 10^−4^, FDR 7.5%) and exhibits genotype-sex interaction effects on *PON2* expression (beta = 0.13; *P*-value = 8.2 × 10^−3^). If tested separately in each sex, this gene-SNP pair is an eQTL in males (*P*-value = 5.5 × 10^−3^) but not females (*P*-value = 5.0 × 10^−1^). Interestingly, *PON2* does not exhibit sex-specific gene expression (*P*-value = 6.3 × 10^−1^, FDR 89.2%) but has been associated with sex-specific effects in oxidative stress responses (Cheng and Klaassen 2012; Giordano et al. 2013; Polonikov et al. 2014).

### Integration of sex-specifics effects with human disease

In humans, sexual dimorphism is observed in the incidence rates and severity of many common diseases, including cardiovascular, immune, and neurological diseases. For example, a recent study (Myers et al. 2014) identified six sex-specific asthma risk loci (*P*-value < 1 × 10^−6^), three of which exhibit genotype-sex interactions on expression in our study (nominal *P*-value < 0.05). Unfortunately, previous limitations on statistical power and study design have provided challenges for identifying significant genotype-sex effects in many disease association studies (Brookes et al. 2004; Wang et al. 2008). Given the role of regulatory variation in determining disease risk (Nica et al. 2010; Nicolae et al. 2010), we asked if genetic variants identified through genome-wide association studies (GWAS) have cumulatively different eQTL effects in males and females. To test this, we obtained trait-associated variants from Immunobase and examined the effect size of each GWAS variant on expression in males and females separately. We found that for the majority of autoimmune diseases, the disease-associated loci exhibit a bias in effect size in one sex (Fig. 5A; see Methods). For example, a significant proportion of genetic variants associated with multiple sclerosis, a disease with well-documented sex-specific disparity in prevalence and severity (Whitacre et al. 1999; Whitacre 2001), exhibit a bias towards females (*P*-value = 5.6 × 10^−7^, binomial exact test). We performed an identical analysis using data available from the NHGRI-EBI GWAS catalog (Welter et al. 2014) and observe similar patterns (Fig. S9). These observations indicate that disease-associated variants have different cumulative effects on gene expression in males and females underlying potential sex-biases in disease prevalence and severity.

**Figure 5.**
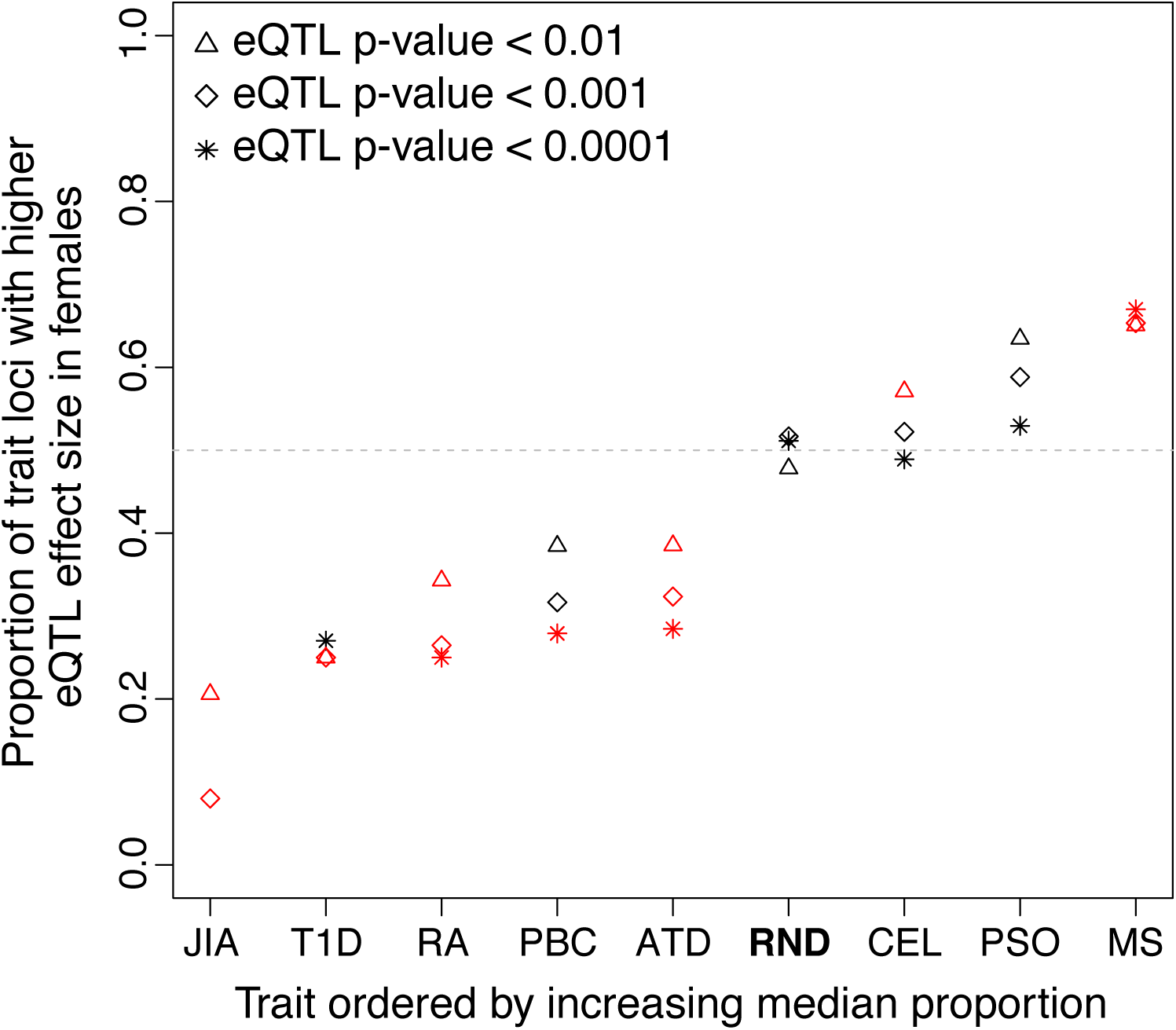
Disease associated variants with sex-biased eQTLs. Proportion of independent (LD-pruned) variants with higher eQTL effect sizes in females for GWAS variants of traits in Immunobase. Tested variants were included based on increasing stringency that they were an eQTL. Red indicates traits with effect sizes that are significantly different between sexes. ATD, autoimmune thyroid disease; CEL, celiac disease, JIA, juvenile idiopathic arthritis; MS, multiple sclerosis; PBC, primarily biliary cirrhosis, PSO, psoriasis, RA, rheumatoid arthritis; RND, random eQTL variants; SLE, systemic lupus erythematous; T1D, type 1 diabetes.

## Discussion

In this study, we have evaluated the effect of the X chromosome and sex on regulatory variation. First, we demonstrated that genes on the X chromosome are more likely to have sex-specific expression compared to genes on the autosomes, and genes with higher sex-specific expression variance are likely to be involved in apoptosis and regulation of cell death. Second, we observe a depletion of regulatory variation on the X chromosome suggesting more efficient purifying selection on the X chromosome relatives to autosomes. In contrast, we tested the effect of genotype-sex interactions across the transcriptome and observed an enrichment of sex-interacting eQTLs on the X chromosome. Third, we investigated the mechanism of sex-interacting expression and eQTLs by generating chromatin accessibility data and demonstrating an enrichment of variants with genotype-sex interactions in regions with differential chromatin accessibility between the sexes. Lastly, we discover that complex traits may have variants with cumulatively different effects between sexes leading to potential sex-biases in their disease prevalence and severity. Together, this work advances our understanding of how the X chromosome and sex shape human gene regulation and highlights their importance in both functional genomic and disease studies.

## Methods

### Study cohort for expression analysis

In the discovery of associations between genotype, expression and sex, we used the Depression, Genes, and Networks (DGN) cohort (Battle et al. 2014; Mostafavi et al. 2014). The cohort is comprised of 922 individuals (648 females and 274 males) of European ancestry between the ages of 21 and 60 years old within the United States. A detailed description of the recruitment and phenotype data for this cohort is provided elsewhere (Battle et al. 2014; Mostafavi et al. 2014).

### Genotype data and imputation

Each individual in the DGN cohort was genotyped for 737,187 single nucleotide polymorphisms (SNPs) located on the autosomes and X chromosome on the Illumina HumanOmni1-Quad BeadChip. Quality control was performed to identify samples with elevated heterozygosity, unexpected ancestry or pairwise IBD, and potential mislabeling, as described elsewhere (Battle et al. 2014). In order to increase the power of the association analyses, we imputed variants onto the genotyped variants. Prior to imputation, we pre-phased the genotypes using SHAPEIT v2.r644 (Delaneau et al. 2012) with default parameters for the autosomes and the ‘--chrX’ flag for the X chromosome. Next, we passed the haplotype estimates to IMPUTE2 (Howie et al. 2009) version 2.2.2 for imputation from the 1000 Genomes Phase I integrated haplotypes (Dec 2013 release) using the following parameters: ‘-Ne 20000 -filt_rules_l eur.maf<0.05 -call_thresh 0.9’. Due to the hemizygosity of males on the X-chromosome, the ‘-chrX’ flag was used in IMPUTE2 to impute the non-pseudo autosomal regions (PAR) of the X chromosome. Quality control after imputation included the exclusion of variants with genotype call rate of <90%, Hardy-Weinberg equilibrium *P*-value of < 1 × 10^−6^, minor allele frequency (MAF) of < 1%, or minimum reference and non-reference allele count (MAC) of 18. A total of 8,465,160 autosomal variants and 156,848 X-chromosome variants passed quality control criteria and were included in the association analyses. Of these, the autosomal variants included 7,714,879 (91.1%) SNPs and 750,281 (8.9%) short insertion-deletion polymorphisms (indels), and the X-chromosome variants included 140,611 (89.6%) SNPs and 16,237 (10.4%) indels.

### Transcriptome data

RNA-sequencing (RNA-seq) was previously performed for each individual in the DGN cohort, as described elsewhere (Battle et al. 2014). In brief, RNA was removed from whole blood, depleted of globin mRNA transcripts, and sequenced in an Illumina HiSeq 200, yielding an average of 70 million 51-bp single-ended reads per individual. Reads were aligned to the NCBI v37 H. sapiens reference genome using TopHat (Trapnell et al. 2009). Gene-level expression was quantified using HTSeq (Anders et al. 2015) and isoform-level expression was quantified using Cufflinks (Roberts et al. 2011). Only uniquely aligned reads were used for expression quantification. We consider a gene expressed if it has at least 10 reads in 100 individuals. Differential expression analysis was conducted using a negative binomial model in the R package DESeq (Anders and Huber 2010) to identify genes with sex-specific expression (FDR 5%) in a matched number of males and females.

### Correction for hidden covariates in transcriptome data

We used the probabilistic estimation of expression residual (PEER) method (Stegle et al. 2012) to estimate and correct for hidden covariates in the RNA-seq data. To infer PEER factors, we supplied the one known covariate (sex) and inferred up to 40 hidden covariates. We calculated the expression residuals from the inferred PEER factors excluding the sex covariate in order to preserve the effect of sex on expression. The number of PEER factors removed was tuned to maximize the number of *cis*-eQTLs in the expression data (Fig. S10). We observed that the removal of these 25 factors from the expression data maximized discoveries.

### Expression variance analysis

To detect genes with sex-specific expression variance, we regressed out known technical factors previously identified in the DGN cohort (Battle et al. 2014). After removing the technical factors, we compared the variance of gene expression across the genome within males and females using the F-test statistic. Specifically, we asked if the expression variance of each gene on the autosomes and X chromosome was significantly different for males and females. We restricted our test to genes with at least ten reads in twenty-five individuals with each sex and matched the number of males and females tested (n = 274). To control for multiple testing, we adjusted the nominal *P*-values using the Bonferroni method and considered genes with adjusted *P*-values < 0.01 as significant. If we altered the method for controlling for multiple testing (e.g. Benjamini-Hochberg or Bonferroni) or the *P*-value threshold for significance, we observed no change to the patterns of sex-specific expression variance across the genome (Fig. 1A). The discoveries within the complete DGN cohort replicated within DGN controls only and within expression data from the ImmVar cohort for CD4 and CD14 cells available from the Gene Expression Omnibus (GEO) under accessible number GSE56035.

### Proportion of variance explained

To determine if higher expression variance on the X chromosome in males was explained by the hemizygous state, we modeled the effect of genotype on expression of genes on the X chromosome in males and females using the following linear model:

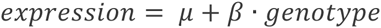

where *μ* is the mean effect and *β* is the effect of genotype on expression. Using an equal number of males and females (*n* = 274), we computed the proportion of variance explained (PVE) by genotype as:

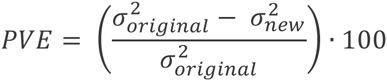

where 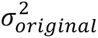 is the variance before we regress out the effect of genotype (given by 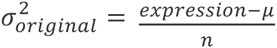 and 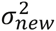 is the variance after we regress out the effect of genotype (given by 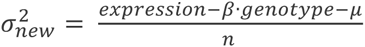).

### ***cis*-QTL mapping.**

We performed association testing for expression QTL (eQTL) using the R package Matrix eQTL (Shabalin 2012). Variants within 1 Mb upstream of the transcription start site (TSS) and 1 Mb downstream of the transcription end site (TES) of a gene were tested using a linear regression model and accounting for sex as a covariate. We model the association between a candidate variant and gene expression by a linear regression:

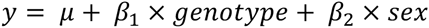

where *y* denotes the observed expression level of the gene, *μ* the mean expression level across the population, *β*_1_ the regression coefficient of genotype, and *β*_2_ the regression coefficient of sex. In the primary QTL analysis, we only tested variants with MAF ≥ 0.05. In the low-frequency QTL analysis, we tested variants with MAF in the range 0.01 to 0.05. To control for multiple testing, we used Bonferroni correction to account for the number of variants tested per gene, retained the best association per gene, and controlled for FDR at the gene-level significance by Benjamini-Hochberg method (Benjamini and Hochberg 1995).

### eQTL effect size

We estimated the eQTL effect sizes by computing fold change, using a subsampling approach to avoid biases from allele frequency. Specifically, we subsampled 100 individuals with an equal distribution of males and females and matched the number of individuals with each allele analyzed. To examine the relation of eQTL effect size and purifying selection ratios, we obtained the ratio of non-synonymous to synonymous substitutions (dN/dS) for human-rhesus gene alignments from Biomart Ensembl (Cunningham et al. 2015).

### Identification of sex-interacting QTLs

We tested for genotype-by-sex (GxS) interactions using the interaction model in the R package Matrix eQTL (Shabalin 2012). This model was used to test for equality of effect sizes between males and females. We model the interaction between genotype and sex by adding an interaction term to the linear regression:

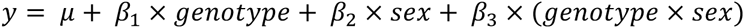

where *y* denotes the observed expression level of the gene, *μ* the mean expression level across the population, *β*_1_ the regression coefficient of genotype, *β*_2_ the regression coefficient of sex, and β_3_ the regression coefficient of the interaction between genotype and sex. For each gene, variants within 1 Mb upstream of the transcription start site (TSS) and 1 Mb downstream of the transcription end site (TES) of a gene were tested. We required variants to have a MAF ≥ 0.05 in the male and female populations. To remove cases where genotype is collinear with sex, we removed variants with a rank less than three between genotype and sex. In addition, we applied a conditional model to improve our statistical power that uses prior regulatory information to detect sex-interacting eQTLs. Specifically, we tested the genotype-sex interaction term only for genetic variants that were previously detected as *cis*-eQTLs in the joint population (adjusted *P*-value < 0.2 after Bonferroni correcting for the number of variants tested per gene). In the conditional model, we improve our power (mean and median improvement of 2.8-fold), but the total number of genes with a sex-interacting eQTL discovered at FDR 5% was unaffected. Therefore, downstream analyses of sex-interacting eQTLs used results from the original approach. To compare the distribution of associations on the autosomes and the X chromosome, we randomly sampled 10,000 gene-variant tests from each distribution (e.g. autosomes and X chromosome) a total of 10,000 times, and calculated the median and 95% confidence intervals for each distribution to create a quantile-quantile (Q-Q) plot (Fig. 3A). All sex-interacting eQTLs tested in the DGN cohort can be visualized at http://montgomerylab.stanford.edu/resources.html.

### Gene ontology enrichment analysis

We used the database for annotation, visualization and integrated discovery (DAVID) (Dennis et al. 2003; Huang da et al. 2009) pathway enrichment online tool to obtain functional annotation for the significant genes in the differential expression variance (see Expression variance analysis) and sex-interacting eQTL (see Identification of sex-interacting eQTLs) analyses. Given that we were interested in the biological processes affecting expression variance and genotype-by-sex interactions, we restricted our gene ontology (GO) analysis to the category biological processes (GOTERM_BP_FA) in DAVID tools. For each GO term category, we used a conservative modified Fisher’s exact test (EASE score in DAVID tools) to test for enrichment against a background gene list and accounted for multiple testing using the Benjamini-Hochberg method (Benjamini and Hochberg 1995). The background gene list consisted of all genes expressed in whole blood (i.e. at least 10 reads in 100 individuals in the DGN cohort).

### Sample collection for open chromatin profiling

We obtained buffy coat samples from twenty healthy donors from the Stanford Blood Center (Stanford, CA). Male and female donors were equally represented for downstream differential analyses. To control for age and ethnicity, the samples selected were restricted to Caucasians between 18 and 45 years old. Age, ethnicity, and healthy status were self-reported by donors.

### PBMC isolation

Peripheral blood mononuclear cells (PBMCs) were isolated from the buffy coats by density gradient centrifugation on using the Ficoll-Paque PLUS (GE Healthcare, Uppsala, Sweden). Briefly, 4 ml of blood was diluted at a ratio of 1:1 (vol/vol) with phosphate-buffered saline (PBS) at room temperature. The 8-ml diluted blood sample was carefully layered onto 6 ml of Ficoll-Pague PLUS in 50-mL conical tubes and centrifuged at 400×g for 30 min at 18°C with the brake off. Next, the lymphocyte layer at the interphase was carefully removed and resuspended in three volumes (6 mL) of PBS, and centrifuged without the brake at 100×g for 10 min at 4°C. To remove platelets from the lymphocyte layer, the cells were resuspended in 6 mL of PBS and centrifuged using the previous settings. Following removal of the supernatant, we counted the cells with a hemocytometer. A subset of 50,000 cells was immediately used for ATAC-seq and the remaining cells were pelleted and stored at -20°C.

### Open chromatin accessibility assay

To profile for open chromatin regions, we used the Assay for Transposase Accessible Chromatin (ATAC-seq) protocol (Buenrostro et al. 2013). The ATAC-seq protocol was followed with zero modifications as previously described (Buenrostro et al. 2015). A total of twenty ATAC-seq libraries with unique single-index barcodes were created for multiplexed sequencing. Each sample was amplified in the range of five to eleven total PCR cycles, as determined by qPCR side reactions performed on each sample. The final amplified libraries were purified using Qiagen MinElute PCR Purification kit and eluted in 20μl elution buffer (10 mM Tris·Cl, pH 8). The fragment size distribution and quality of the libraries were evaluated on an Agilent Bioanalyzer 2000. Each library was quantified using the KAPA Library Quantification Kit (KAPA Biosystems) in triplicate at two dilutions, 1:10^5^ and 1:10^6^. Based on the individual library concentrations, the libraries were pooled at an equimolar concentration for 76-bp paired-end sequencing on an Illumina NextSeq 500 platform.

### Open chromatin accessibility data processing and peak calling

We obtained approximately 200 million paired-end 76-bp reads from pooled sequencing and de-multiplexed the reads as BCL files were converted to FASTQ files using the Illumina BCL2FASTQ Conversion Software (version 2.15.0). Reads were aligned to hg19 using Bowtie2 (Langmead and Salzberg 2012). Reads were filtered by removing those with an alignment quality score less than 4 and reads mapping to chromosome Y, mitochondria, or unmapped contigs.

For comparison between samples, we obtained a consensus set of PBMC ATAC-seq regions by calling peaks on the pooled reads with MAC2 (Zhang et al. 2008) using the following parameters: “-g hs --q 0.01 --nomodel --nolambda --keep-dup all --call-summits.” The peaks were filtered using consensus excludable ENCODE blacklist to remove artificial signal in certain regions of the genome, such as centromeres, telomeres, and simple repeats (Consortium 2012). For quantification of peaks, peak bed files were converted to GFF3 format and the number of reads per peak per sample was quantified with HTSeq (Anders and Huber 2010) using the following parameters: “htseq-count -m intersection-strict --stranded=no’. We retained peaks that contain at least twenty total reads and read coverage across at least four individuals for downstream analysis.

### Differential open chromatin accessibility analysis

To identify regions with differential chromatin accessibility between males and females, we used the R package DESeq (Anders and Huber 2010). The data were normalized by the effective library size and variance was estimated for each group. We tested for differential chromatin accessibility using a model based on the negative binomial distribution. To account for multiple testing, we adjusted the *P*-values by the Benjamini-Hochberg method; regions with a FDR < 0.05 were considered to be regions with differential open chromatin.

### Trait Analysis

We obtained trait-associated SNPs from Immunobase (available at <u>www.immunobase.org</u>; accessed on April 2, 2015) and the NHGRI-EBI GWAS catalog (Welter et al. 2014) (available at http://www.ebi.ac.uk/gwas; accessed on April 2, 2015). We only considered independent variants (R^2^ < 0.5) with a genome-wide association threshold of *P*-value < 1 × 10^−8^. In the DGN cohort, we calculated the absolute effect size of each independent GWAS variant on expression for genes within 1 Mb in the male (n = 274) and subsampled female (n = 274) populations. We considered a GWAS variant to have a sex-specific effect if the effect size in one sex was 1.2-fold greater than the opposite sex. For variants with a 1.2 fold greater effect in either sex, we ran a two-sided binomial exact test to determine if one sex had greater effects than the opposite sex. Only traits with at least twenty significant GWAS variants were considered in the trait analysis (Fig. 5A).

## Data Access

Genotype, raw RNA-seq, quantified expression, and covariate data for the DGN cohort are available by application through the NIMH Center for Collaborative Genomic Studies on Mental Disorders. Instructions for requesting access to data can be found at <u>www.nimhgenetics.org/access_data_biomaterial.php,</u> and inquiries should reference the “Depression Genes and Networks study (D. Levinson, PI)". Open chromatin accessibility data is available on the Gene Expression Omnibus (GEO) under accessible number GSE69749.

## Acknowledgments

We would like to thank Stephen J. Galli for critical review of the manuscript. KRK is supported by DoD, Air Force Office of Scientific Research, National Defense Science and Engineering Graduate (NDSEQ) Fellowship, 32 CFR 168a. SBM is supported by the Edward Mallinckrodt Jr. Foundation. AJB and SBM are supported by the NIH (R01MH101814).

## References

Anders S, Huber W. 2010. Differential expression analysis for sequence count data. Genome Biol 11(10): R106.

Anders S, Pyl PT, Huber W. 2015. HTSeq-a Python framework to work with high-throughput sequencing data. Bioinformatics 31(2): 166–169.

Andolfatto P. 2001. Contrasting patterns of X-linked and autosomal nucleotide variation in Drosophila melanogaster and Drosophila simulans. Mol Biol Evol 18(3): 279–290.

Awadalla P, Boileau C, Payette Y, Idaghdour Y, Goulet JP, Knoppers B, Hamet P, Laberge C, Project CA. 2013. Cohort profile of the CARTaGENE study: Quebec’s population-based biobank for public health and personalized genomics. International journal of epidemiology 42(5): 1285–1299.

Battle A, Mostafavi S, Zhu X, Potash JB, Weissman MM, McCormick C, Haudenschild CD, Beckman KB, Shi J, Mei R et al. 2014. Characterizing the genetic basis of transcriptome diversity through RNA-sequencing of 922 individuals. Genome Res 24(1): 14–24.

Benjamini Y, Hochberg Y. 1995. Controlling the False Discovery Rate - a Practical and Powerful Approach to Multiple Testing. J Roy Stat Soc B Met 57(1): 289–300.

Breslau N, Davis GC, Andreski P, Peterson EL, Schultz LR. 1997. Sex differences in posttraumatic stress disorder. Archives of general psychiatry 54(11): 1044–1048.

Brookes ST, Whitely E, Egger M, Smith GD, Mulheran PA, Peters TJ. 2004. Subgroup analyses in randomized trials: risks of subgroup-specific analyses; power and sample size for the interaction test. Journal of clinical epidemiology 57(3): 229–236.

Buenrostro JD, Giresi PG, Zaba LC, Chang HY, Greenleaf WJ. 2013. Transposition of native chromatin for fast and sensitive epigenomic profiling of open chromatin, DNA-binding proteins and nucleosome position. Nat Methods 10(12): 1213–1218.

Buenrostro JD, Wu B, Chang HY, Greenleaf WJ. 2015. ATAC-seq: A Method for Assaying Chromatin Accessibility Genome-Wide. Current protocols in molecular biology / edited by Frederick M Ausubel [et al] 109: 2129 21–29.

Carrel L, Willard HF. 2005. X-inactivation profile reveals extensive variability in X-linked gene expression in females. Nature 434(7031): 400–404.

Chang D, Gao F, Slavney A, Ma L, Waldman YY, Sams AJ, Billing-Ross P, Madar A, Spritz R, Keinan A. 2014. Accounting for eXentricities: analysis of the X chromosome in GWAS reveals X-linked genes implicated in autoimmune diseases. PLoS One 9(12): e113684.

Cheng X, Klaassen CD. 2012. Hormonal and chemical regulation of paraoxonases in mice. J Pharmacol Exp Ther 342(3): 688–695.

Cohn BA, Wingard DL, Cirillo PM, Cohen RD, Reynolds P, Kaplan GA. 1996. Differences in lung cancer risk between men and women: Examination of the evidence. J Natl Cancer I 88(24): 1867–1867.

Consortium EP. 2012. An integrated encyclopedia of DNA elements in the human genome. Nature 489(7414): 57–74.

Consortium GT. 2013. The Genotype-Tissue Expression (GTEx) project. Nat Genet 45(6): 580–585.

Consortium GT. 2015. Human genomics. The Genotype-Tissue Expression (GTEx) pilot analysis: multitissue gene regulation in humans. Science 348(6235): 648–660.

Cunningham F, Amode MR, Barrell D, Beal K, Billis K, Brent S, Carvalho-Silva D, Clapham P, Coates G, Fitzgerald S et al. 2015. Ensembl 2015. Nucleic Acids Res 43(Database issue): D662–669.

Delaneau O, Marchini J, Zagury JF. 2012. A linear complexity phasing method for thousands of genomes. Nat Methods 9(2): 179–181.

Dennis G, Jr., Sherman BT, Hosack DA, Yang J, Gao W, Lane HC, Lempicki RA. 2003. DAVID: Database for Annotation, Visualization, and Integrated Discovery. Genome Biol 4(5): P3.

Dimas AS, Deutsch S, Stranger BE, Montgomery SB, Borel C, Attar-Cohen H, Ingle C, Beazley C, Gutierrez Arcelus M, Sekowska M et al. 2009. Common regulatory variation impacts gene expression in a cell type-dependent manner. Science 325(5945): 1246–1250.

Dimas AS, Nica AC, Montgomery SB, Stranger BE, Raj T, Buil A, Giger T, Lappalainen T, Gutierrez-Arcelus M, McCarthy MI et al. 2012a. Sex-biased genetic effects on gene regulation in humans. Genome Research 22(12): 2368–2375.

Dimas AS, Nica AC, Montgomery SB, Stranger BE, Raj T, Buil A, Giger T, Lappalainen T, Gutierrez-Arcelus M, Mu TC et al. 2012b. Sex-biased genetic effects on gene regulation in humans. Genome Res 22(12): 2368–2375.

Dobyns WB, Filauro A, Tomson BN, Chan AS, Ho AW, Ting NT, Oosterwijk JC, Ober C. 2004. Inheritance of most X-linked traits is not dominant or recessive, just X-linked. American journal of medical genetics Part A 129A(2): 136–143.

Giordano G, Tait L, Furlong CE, Cole TB, Kavanagh TJ, Costa LG. 2013. Gender differences in brain susceptibility to oxidative stress are mediated by levels of paraoxonase-2 expression. Free radical biology & medicine 58: 98–108.

Hankin BL, Abramson LY. 2001. Development of gender differences in depression: an elaborated cognitive vulnerability-transactional stress theory. Psychological bulletin 127(6): 773–796.

Heid IM Jackson AU Randall JC Winkler TW Qi L Steinthorsdottir V Thorleifsson G Zillikens MC Speliotes EK Magi R et al. 2010. Meta-analysis identifies 13 new loci associated with waist-hip ratio and reveals sexual dimorphism in the genetic basis of fat distribution. Nat Genet 42(11): 949–960.

Howie BN, Donnelly P, Marchini J. 2009. A flexible and accurate genotype imputation method for the next generation of genome-wide association studies. PLoS Genet 5(6): e1000529.

Huang da W, Sherman BT, Lempicki RA. 2009. Systematic and integrative analysis of large gene lists using DAVID bioinformatics resources. Nature protocols 4(1): 44–57.

Hussin JG, Hodgkinson A, Idaghdour Y, Grenier JC, Goulet JP, Gbeha E, Hip-Ki E, Awadalla P. 2015. Recombination affects accumulation of damaging and disease-associated mutations in human populations. Nat Genet 47(4): 400–404.

Idaghdour Y, Awadalla P. 2012. Exploiting gene expression variation to capture gene-environment interactions for disease. Frontiers in genetics 3: 228.

Lang JT, McCullough LD. 2008. Pathways to ischemic neuronal cell death: are sex differences relevant? Journal of translational medicine 6: 33.

Langmead B, Salzberg SL. 2012. Fast gapped-read alignment with Bowtie 2. Nat Methods 9(4): 357–359.

Lappalainen T, Sammeth M, Friedlander MR, Hoen PA, Monlong J, Rivas MA, Gonzalez-Porta M, Kurbatova N, Griebel T, Ferreira PG et al. 2013. Transcriptome and genome sequencing uncovers functional variation in humans. Nature 501(7468): 506–511.

Lerner DJ, Kannel WB. 1986. Patterns of coronary heart disease morbidity and mortality in the sexes: a 26-year follow-up of the Framingham population. American heart journal 111(2): 383–390.

Liu CT, Estrada K, Yerges-Armstrong LM, Amin N, Evangelou E, Li G, Minster RL, Carless MA, Kammerer CM, Oei L et al. 2012a. Assessment of gene-by-sex interaction effect on bone mineral density. Journal of bone and mineral research: the official journal of the American Society for Bone and Mineral Research 27(10): 2051–2064.

Liu LY, Schaub MA, Sirota M, Butte AJ. 2012b. Sex differences in disease risk from reported genome-wide association study findings. Hum Genet 131(3): 353–364.

Lu J, Wu CI. 2005. Weak selection revealed by the whole-genome comparison of the X chromosome and autosomes of human and chimpanzee. Proc Natl Acad Sci U S A 102(11): 4063–4067.

Luan JA, Wong MY, Day NE, Wareham NJ. 2001. Sample size determination for studies of gene-environment interaction. International journal of epidemiology 30(5): 1035–1040.

Mendelsohn ME, Karas RH. 2005. Molecular and cellular basis of cardiovascular gender differences. Science 308(5728): 1583–1587.

Molloy EJ, O’Neill AJ, Grantham JJ, Sheridan-Pereira M, Fitzpatrick JM, Webb DW, Watson RW. 2003. Sex-specific alterations in neutrophil apoptosis: the role of estradiol and progesterone. Blood 102(7): 2653–2659.

Montgomery SB, Sammeth M, Gutierrez-Arcelus M, Lach RP, Ingle C, Nisbett J, Guigo R, Dermitzakis ET. 2010. Transcriptome genetics using second generation sequencing in a Caucasian population. Nature 464(7289): 773–777.

Mostafavi S, Battle A, Zhu X, Potash JB, Weissman MM, Shi J, Beckman K, Haudenschild C, McCormick C, Mei R et al. 2014. Type I interferon signaling genes in recurrent major depression: increased expression detected by whole-blood RNA sequencing. Molecular psychiatry 19(12): 1267–1274.

Myers RA, Scott NM, Gauderman WJ, Qiu W, Mathias RA, Romieu I, Levin AM, Pino-Yanes M, Graves PE, Villarreal AB et al. 2014. Genome-wide interaction studies reveal sex-specific asthma risk alleles. Hum Mol Genet 23(19): 5251–5259.

Naugler WE, Sakurai T, Kim S, Maeda S, Kim K, Elsharkawy AM, Karin M. 2007. Gender disparity in liver cancer due to sex differences in MyD88-dependent IL-6 production. Science 317(5834): 121–124.

Nica AC, Montgomery SB, Dimas AS, Stranger BE, Beazley C, Barroso I, Dermitzakis ET. 2010. Candidate causal regulatory effects by integration of expression QTLs with complex trait genetic associations. PLoS Genet 6(4): e1000895.

Nicolae DL, Gamazon E, Zhang W, Duan SW, Dolan ME, Cox NJ. 2010. Trait-Associated SNPs Are More Likely to Be eQTLs: Annotation to Enhance Discovery from GWAS. Plos Genetics 6(4).

Ober C, Loisel DA, Gilad Y. 2008. Sex-specific genetic architecture of human disease. Nat Rev Genet 9(12): 911–922.

Park C, Carrel L, Makova KD. 2010. Strong Purifying Selection at Genes Escaping X Chromosome Inactivation. Molecular Biology and Evolution 27(11): 2446–2450.

Patsopoulos NA, Tatsioni A, Ioannidis JP. 2007. Claims of sex differences: an empirical assessment in genetic associations. JAMA: the journal of the American Medical Association 298(8): 880–893.

Pickrell JK, Marioni JC, Pai AA, Degner JF, Engelhardt BE, Nkadori E, Veyrieras JB, Stephens M, Gilad Y, Pritchard JK. 2010. Understanding mechanisms underlying human gene expression variation with RNA sequencing. Nature 464(7289): 768–772.

Pigott TA. 1999. Gender differences in the epidemiology and treatment of anxiety disorders. The Journal of clinical psychiatry 60 Suppl 18: 4–15.

Polonikov AV, Ivanov VP, Bogomazov AD, Freidin MB, Illig T, Solodilova MA. 2014. Antioxidant defense enzyme genes and asthma susceptibility: gender-specific effects and heterogeneity in gene-gene interactions between pathogenetic variants of the disease. BioMed research international 2014: 708903.

Randall JC Winkler TW Kutalik Z Berndt SI Jackson AU Monda KL Kilpelainen TO Esko T Magi R Li S et al. 2013. Sex-stratified genome-wide association studies including 270,000 individuals show sexual dimorphism in genetic loci for anthropometric traits. PLoS Genet 9(6): e1003500.

Ranz JM, Castillo-Davis CI, Meiklejohn CD, Hartl DL. 2003. Sex-dependent gene expression and evolution of the Drosophila transcriptome. Science 300(5626): 1742–1745.

Rice WR. 1984. Sex Chromosomes and the Evolution of Sexual Dimorphism. Evolution 38(4): 735–742.

Roberts A, Trapnell C, Donaghey J, Rinn JL, Pachter L. 2011. Improving RNA-Seq expression estimates by correcting for fragment bias. Genome Biol 12(3): R22.

Schaffner SF. 2004. The X chromosome in population genetics. Nat Rev Genet 5(1): 43-51.

Shabalin AA. 2012. Matrix eQTL: ultra fast eQTL analysis via large matrix operations. Bioinformatics 28(10): 1353–1358.

Stegle O, Parts L, Piipari M, Winn J, Durbin R. 2012. Using probabilistic estimation of expression residuals (PEER) to obtain increased power and interpretability of gene expression analyses. Nature protocols 7(3): 500–507.

Trapnell C, Pachter L, Salzberg SL. 2009. TopHat: discovering splice junctions with RNA-Seq. Bioinformatics (Oxford, England) 25(9): 1105–1111.

Tukiainen T, Pirinen M, Sarin AP, Ladenvall C, Kettunen J, Lehtimaki T, Lokki ML, Perola M, Sinisalo J, Vlachopoulou E et al. 2014. Chromosome X-wide association study identifies Loci for fasting insulin and height and evidence for incomplete dosage compensation. PLoS Genet 10(2): e1004127.

Vicoso B, Charlesworth B. 2006. Evolution on the X chromosome: unusual patterns and processes. Nat Rev Genet 7(8): 645–653.

Wang C, Cheng Y, Liu T, Li Q, Fillingim RB, Wallace MR, Staud R, Kaplan L, Wu R. 2008. A computational model for sex-specific genetic architecture of complex traits in humans: implications for mapping pain sensitivity. Molecular pain 4: 13.

Welter D, MacArthur J, Morales J, Burdett T, Hall P, Junkins H, Klemm A, Flicek P, Manolio T, Hindorff L et al. 2014. The NHGRI GWAS Catalog, a curated resource of SNP-trait associations. Nucleic Acids Res 42(Database issue): D1001–1006.

Whitacre CC. 2001. Sex differences in autoimmune disease. Nat Immunol 2(9): 777–780.

Whitacre CC, Reingold SC, O’Looney PA. 1999. A gender gap in autoimmunity. Science 283(5406): 1277–1278.

Wise AL, Gyi L, Manolio TA. 2013. eXclusion: toward integrating the X chromosome in genome-wide association analyses. Am J Hum Genet 92(5): 643–647.

Yang X, Schadt EE, Wang S, Wang H, Arnold AP, Ingram-Drake L, Drake TA, Lusis AJ. 2006. Tissue-specific expression and regulation of sexually dimorphic genes in mice. Genome Res 16(8): 995–1004.

Yao C, Joehanes R, Johnson AD, Huan T, Esko T, Ying S, Freedman JE, Murabito J, Lunetta KL, Metspalu A et al. 2014. Sex-and age-interacting eQTLs in human complex diseases. Hum Mol Genet 23(7): 1947–1956.

Ye CJ, Feng T, Kwon HK, Raj T, Wilson MT, Asinovski N, McCabe C, Lee MH, Frohlich I, Paik HI et al. 2014. Intersection of population variation and autoimmunity genetics in human T cell activation. Science 345(6202): 1254665.

Zhang Y, Liu T, Meyer CA, Eeckhoute J, Johnson DS, Bernstein BE, Nusbaum C, Myers RM, Brown M, Li W et al. 2008. Model-based analysis of ChIP-Seq (MACS). Genome Biol 9(9): R137.

